# Fast and Accurate Prediction of Intrinsically Disordered Protein by Protein Language Model

**DOI:** 10.1101/2022.10.15.512345

**Authors:** Shijie Xu, Akira Onoda

## Abstract

**Motivation:** Intrinsically disordered proteins (IDPs) play a vital role in various biological processes and have attracted increasing attention in the last decades. Predicting IDPs from primary structures of proteins provides a very useful tool for protein analysis. However, most of the existing prediction methods heavily rely on multiple sequence alignments (MSAs) of homologous sequences which are formed by evolution over billions of years. Obtaining such information requires searching against the whole protein databases to find similar sequences and since this process becomes increasingly time-consuming, especially in large-scale practical applications, the alternative method is needed.

**Results:** In this paper, we proposed a novel IDP prediction method named IDP-PLM, based on the protein language model (PLM). The method does not rely on MSAs or MSA-based profiles but leverages only the protein sequences, thereby achieving state-of-the-art performance even compared with predictors using protein profiles. The proposed IDP-PLM is composed of stacked predictors designed for several different protein-related tasks: secondary structure prediction, linker prediction, and binding predictions. In addition, predictors for the single task also achieved the highest accuracy. All these are based on PLMs thus making IDP-PLM not rely on MSA-based profiles. The ablation study reveals that all these stacked predictors contribute positively to the IDP prediction performance of IDP-PLM.

**Availability:** The method is available at http://github.com/xu-shi-jie.

**Supplementary information:** Supplementary data are available at *Bioinformatics* online.

## 1 Introduction

Intrinsically disordered proteins (IDPs) and intrinsically disordered regions (IDRs) are proteins and regions lacking stable structures but play a crucial role in biological processes (Jirgensons, 1958). Recent research has shown their wide occurrence in all domains of life (Peng *et al.*, 2015). For example, more than 6% amino acid residues in annotated sequences exist as disordered in SwissProt (Boeckmann *et al.*, 2003) and this ratio increases even over 30% in eukaryotes (Xue *et al.*, 2012). In addition, quite a few human diseases such as Alzheimer’s disease (Carballo-Pacheco and Strodel, 2017), Parkinson’s disease (Coskuner and Uversky, 2019) and cancers (Rajagopalan *et al.*, 2011) have been indicated to be related to IDRs, which have attracted the interests in the application for drug design (Ambadipudi and Zweckstetter, 2016). Despite the importance and potential applications of IDRs, it is significantly expensive and laborious to identify and annotate them and there are only thousands of IDRs recognized experimentally (Quaglia *et al.*, 2022).

The computational methods that predict IDRs directly from sequences provide very useful tools for protein analysis and numerous methods have been proposed in the past decades. With the help of modern deep learning, neural network-based methods (Wang *et al.*, 2016a; Liu *et al.*, 2018; Tang *et al.*, 2020; Liu *et al.*, 2021; Hu *et al.*, 2021) predominate in the field. They are built on different architectures such as long short-term memory networks (LSTM) (Hochreiter and Schmidhuber, 1997), convolutional networks (LeCun *et al.*, 1998) and self-attention mechanism (Vaswani *et al.*, 2017). To achieve better performance, the majority of existing methods depend largely on protein profiles generated by multiple sequence alignments (MSAs) of homologous sequences, e.g., position-specific scoring matrix (PSSM) (Altschul *et al.*, 1997) and hidden Markov models (HMM) (Eddy, 1998). However, this could bring two problems: 1) MSAs of disordered regions are not as good as those of ordered regions. Compared to ordered regions, the rapid evolution of IDRs (Zarin *et al.*, 2019) makes homologous proteins extraordinarily diverse in sequences. Thus it is burdensome to find similar sequences as well as align them, thereby reducing the performance of predictors. 2) Generating MSAs suffers from the time-consuming searching against the whole protein databases, e.g., UniRef100 containing over 0.3 billion protein sequences (UniProt, 2021). With the rapid increase of sequences in databases, generating MSA-based profiles in a reasonable time becomes increasingly difficult. In addition, other non-MSA profiles have also been proven useful for IDR prediction. For example, the structural information including predicted residue-residue contacts (Seemayer *et al.*, 2014) and predicted secondary structures (Yang *et al.*, 2017) are also utilized (Hanson *et al.*, 2017).

Recently, the large-scale language model (LM) (Devlin *et al.*, 2018) has been arising as a vital method in machine learning. LMs are neural network-based models trained on large-scale unlabeled language corpus from which they can learn the patterns of sequences in a self-supervised way. This procedure is called pre-training and the pre-trained models have shown to be superior to conventional methods in a wide range of machine learning tasks. LMs trained on protein sequences, named protein language models (PLMs) (Elnaggar *et al.*, 2020; Rives *et al.*, 2021), are capable of extracting structural information efficiently yet effectively and reaching competitive results compared to conventional methods on several protein-related tasks including secondary structure prediction (Elnaggar *et al.*, 2020) and residue-residue contact prediction (Rives *et al.*, 2021).

A few works focusing on PLM-based IDR prediction achieved comparable results (Papastratis, 2022; Tamiola *et al.*, 2022). Their architectures are designed by adding extra modules to PLMs, i.e., softmax layer or logistic regression. These shallow modules have no sufficient capacity to store learned IDR-related information. Therefore, their methods can only benefit from PLMs. On the other hand, state-of-the-art conventional machine learning methods for IDR prediction work excellently with ingenious architectures, which often contain a variety of modules responsible for extracting various useful information from the sequences, e.g., predicted contact map and solvent exposure (Hanson *et al.*, 2017). To the best of our knowledge, there is currently no such work demonstrating how to combine the PLMs with conventional methods and further improve the performance of IDR prediction. To exploit the advantages of PLMs such as high efficiency as well as the success of conventional methods in excellent accuracy, a fast yet accurate approach is immediately desired.

In this study, we proposed a novel IDR prediction method named IDP-PLM aiming to predict IDRs based on the PLMs. It consists of several independent modules including a secondary structure predictor, a linker predictor and several binding predictors including protein-, DNA-, RNA- and lipid-binding predictor. To avoid dependence on MSA-based protein profiles, these independent predictors are based on PLM appending with an LSTM network and this makes our IDP-PLM use only the deep learning-based profiles. Compared to the existing work, all of them achieved state-of-the-art performance on the test datasets in terms of all or part of the measures. By stacking them together with extra useful modules into an integrated architecture, the proposed IDP-PLM achieved state-of-the-art performance in terms of both efficiency and accuracy. Moreover, we studied the influences of each module in IDP-PLM by conducting ablation analyses and found that all modules were necessary and contributed positively to the accuracy of IDR prediction.

## 2 Materials and methods

### 2.1 Datasets

#### Secondary structure, protein-, DNA-, RNA-binding, linker and lipid-binding benchmark datasets

For secondary structure prediction, we used the benchmark dataset NetSurfP-2.0 (Klausen *et al.*, 2019) for training and CB513 (Cuff and Barton, 1999) for test, which contain 10,893 and 513 protein sequences with annotated secondary structures respectively. Protein-, DNA- and RNA-binding prediction is performed by the benchmark datasets used in the reported work for training and test which contain 767 and 86 protein sequences with annotations, respectively (Zhang *et al.*, 2022). Linker prediction is performed by the benchmark datasets used in the reported work for training and test which contain 144 and 60 protein sequences with annotations, respectively (Meng and Kurgan, 2016). Lipid-binding prediction is performed by the benchmark datasets used in the reported work for training and test which contain 222 and 252 protein sequences with annotations, respectively (Katuwawala *et al.*, 2021). The detailed statistical information of the linker, protein-, DNA-, RNA- and lipid-binding datasets is summarized in Supplementary Table S1.

#### IDR benchmark datasets

We constructed a dataset for training the IDR predictor by combining three different datasets including DM4229 (Zhang *et al.*, 2012), the training dataset in the reported work (Liu *et al.*, 2018) which is extracted from MobiDB (Piovesan *et al.*, 2021) and DisProt dataset (ver. 2022_06) (Quaglia *et al.*, 2022) which consist of 4,229, 5,273 and 2,416 protein sequences, respectively, and the united dataset contains 11,902 protein sequences after removing redundancy and further processing. To comprehensively evaluate the performance of our method, we tested it on several independent datasets including DISORDER723 (Cheng *et al.*, 2005), MXD494 (Peng and Kurgan, 2012) as well as the very recent CAID (Necci *et al.*, 2021). For a fair comparison, the training dataset of each test was constructed separately from the aforementioned united dataset by removing sequences with the identity greater than 25% against the test dataset through *Blastclust* tool (Altschul *et al.*, 1997). This makes the models cannot see any leaked information from the test datasets during the training. Their detailed statistical information is summarized in Supplementary Fig. S1 and Table S2.

### 2.2 Protein language model

It has been a long-standing problem in bioinformatics that state-of-the-art models focusing on protein function and structure predictions require a variety of protein profiles as inputs which usually need a time-consuming search to generate multiple sequence alignments (MSAs). They are, however, not sufficiently powerful in some cases (e.g., the dark proteome) (Perdigão and Rosa, 2019; Zarin *et al.*, 2019) and cannot differentiate the proteins in the same family. Therefore, non-MSA-based models which do not leverage any protein profiles based on MSAs and can work with sequences are urgent for a long time in the bioinformatics community.

Benefiting from the rapid development of machine learning, a series of efforts addressing this problem have been proposed (Iuchi *et al.*, 2021). The protein language model (PLM) which struggles to extract essential information from oceans of protein sequences has been arising as a potential direction. A PLM receives a protein sequence as the only input and outputs vectors for each amino acid residue in the proteins, which can be utilized for subsequent protein-related tasks, e.g., ProtVecX (Asgari *et al.*, 2019) and SeqVec (Heinzinger *et al.*, 2019) have achieved excellent performance in various protein prediction tasks. Their success without using MSA-generated profiles suggested that protein representations generated by PLMs can play a similar role as MSA-generated profiles and improve the performance of various protein prediction tasks, achieving good performance approximate to conventional models. However, their method did not reach the best performance in terms of accuracy on quite a few tasks, due to the limitation of architecture and model size.

Recently, the large-scale Transformer-based (Vaswani *et al.*, 2017) neural networks built with the masked language model (MLM) (Devlin *et al.*, 2018) have achieved remarkable progress in natural language processing and computer vision. Transformers are large-scale neural networks built with multiple self-attention layers, which have been shown very useful to capture long-distance interactions in the sequence. In MLM, the tokens in the sequence are masked randomly with a specific ratio and the model would try to recover the masked tokens according to the corrupted sequence. The objective is to minimize the following negative log-likelihood function (Rives *et al.*, 2021):

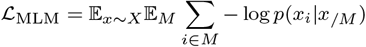

where *X* is the corpus, *M* is the mask, *x_i_* is the *i*^th^ token in the sequence, *x/M* is the unmasked tokens and *p* represents the probability of *x_i_* occurring in the corrupted sequence.

Since protein primary structures are considered as the sequences of amino acid residues, MLM is also applicable to proteins. The latest work dedicated to studying PLM and protein-related problems has shed a light on approaching the goal to surpass the conventional models (Elnaggar *et al.*, 2020; Rives *et al.*, 2021). Although current works resulted in these PLM-based models approaching yet slightly lower than the conventional models on the majority of protein prediction tasks such as residue-residue contact prediction (Rao *et al.*, 2020), secondary structure prediction (Rives *et al.*, 2021) and three-dimensional prediction (Lin *et al.*, 2022), there could be many possibilities to exploit the PLM-based methods for specific tasks such as IDR prediction by employing better architectures.

### 2.3 Long short-term memory network

Long short-term memory network (LSTM) (Hochreiter and Schmidhuber, 1997) is a variant of recurrent neural network (RNN) (Rumelhart *et al.*, 1985) and has been proposed to solve sequential problems in machine learning including human language translation and sequence labeling. In contrast to the vanilla RNNs suffering from the inability of memorizing long-distance information in the sequence, LSTM behaves better in practice through a well-designed gated mechanism that retains the important long-distance information while ignoring the unimportant short-distance information. LSTM holds a prevailing position in protein sequence learning since proteins are essentially the sequences of amino acid residues and even far longer than sentences in human languages, which typically have less than 50 words in English while proteins consist of hundreds of amino acid residues on average. In this work, we adopted the bidirectional LSTM (BiLSTM) network consisting of two LSTM networks by obtaining the information from both two directions of sequences.

### 2.4 Architecture of our method

Based on the above discussions, we proposed our method and its architecture is shown in Figure 1(a). It adopts a protein language model, a secondary structure predictor, a protein-binding predictor, a linker predictor, DNA-, RNA-, lipid-binding predictor, an ADD-MEAN module, a BiLSTM network and a fully connected layer. The arrows represent the data flow, where a batch of protein sequences is fed into the model and the outputs are the predicted intrinsic disorder of the proteins where 1 stands for disorder and 0 for order. The modules with corners mean that they are trainable while others are not, and different colors are used to differentiate them in functions.

**Fig. 1.**
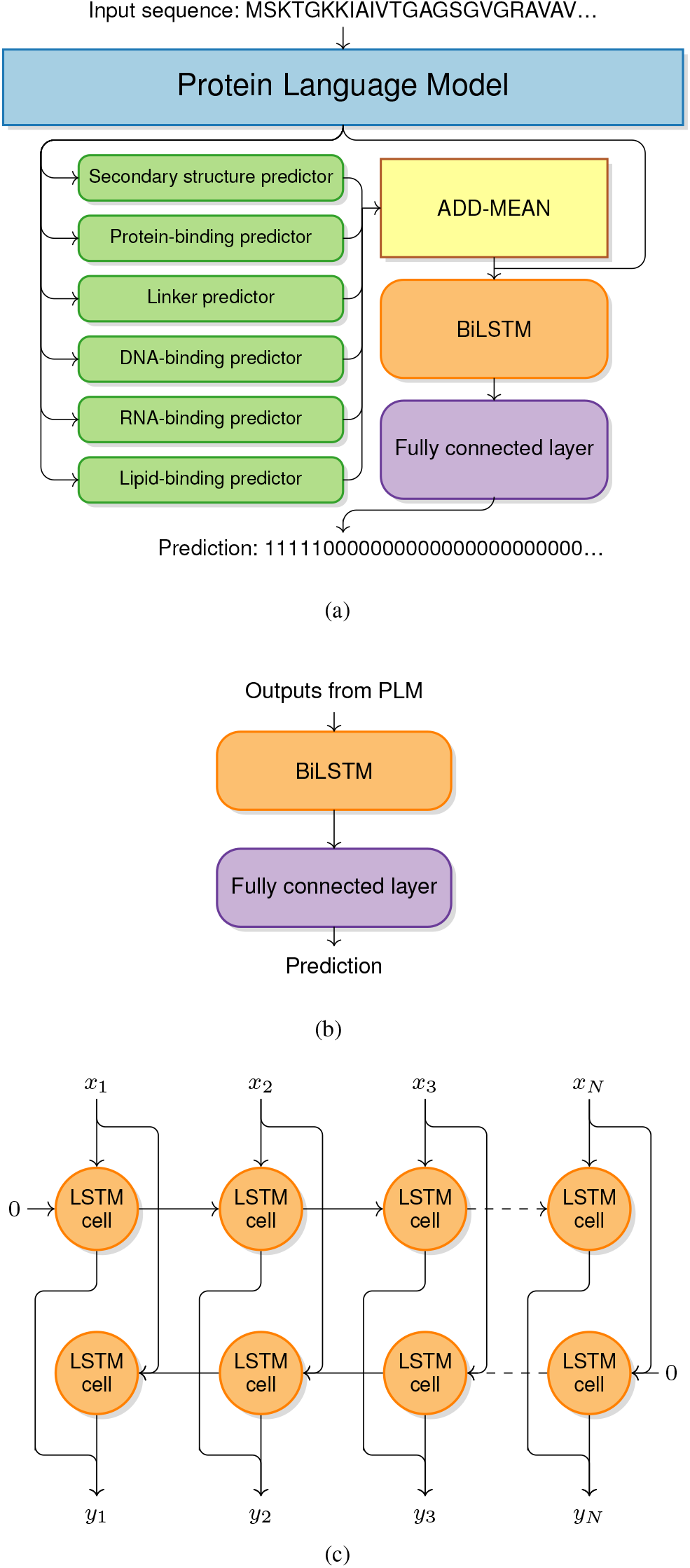
(a) The overall architecture of IDP-PLM predictor. (b) The shared architecture of the secondary structure, linker, protein-, DNA-, RNA- and lipid-binding predictors. (c) The architecture of the BiLSTM network.

#### Protein language model

Since it is tremendously expensive and unnecessary to train a large-scale protein language model from scratch, we employed a publicly available pre-trained model named ESM-1b (Rives *et al.*, 2021), which has 34 layers, 1280-dimensional outputs, and approximately 650M parameters trained on UniRef50 (UniProt, 2021). Due to the limitation of the maximum sequence length of ESM-1b being 1024 which contains two special tokens, i.e., the beginning and end of the sequence, we cut all protein sequences longer than 1022 into the segments within the length of 1022. The parameters of the pre-trained model are frozen during the training for the reason of hardware constraints.

#### Secondary structure predictor, protein-binding predictor, linker predictor, DNA-, RNA-, and lipid-binding predictor

We employed the BiLSTM network as the backbone network for our secondary structure predictor, protein-binding predictor, linker predictor, DNA-, RNA-, and lipid-binding predictor. The outputs from PLM will be fed into a BiLSTM network and a single fully connected layer is employed to convert its output into the final prediction, as shown in Figure 1(b). All of these predictors share a common architecture: three-layer BiLSTM with 1280-dimensional inputs as well as 512 hidden units and a fully connected layer with 8- (for 8-state secondary structure prediction) or 2- (for other prediction) dimensional outputs.

#### ADD-MEAN

Inspired by the work of flDPnn (Hu *et al.*, 2021) which suggested that protein-level information is beneficial for the IDR prediction, we designed an ADD-MEAN module to add environmental information for each amino acid residue. Suppose *e*_1_, *e*_2_, . . . , *e_N_* are the outputs vectors of predictor (e.g. secondary structure predictor) for each amino acid residue in the protein where *N* is the number of residues, and 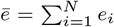 is the averaged representation of the protein, then the ADD-MEAN module would produce the concatenated vectors

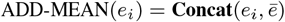

for the *i*^th^ residue where **Concat** means concatenation of vectors.

#### BiLSTM and fully connected layer

The detailed architecture of the BiLSTM network can be seen in Figure 1(c) where the circles mean LSTM cells and the arrows mean the flow of data. For each LSTM cell, it receives an input as well as the previous cell’s states where the initial states are set to zero in our implementation. When the output of the current LSTM cell is used as the input of another LSTM cell and this action is repeated for *N* times, a multi-layer BiLSTM network has been built. We also adopted the fully connected layers appended below the BiLSTM network to transfer outputs into the final results. They are the functions on each residue: *f* (*x*) = *σ*(*Wx* + *b*) where *x* is the representation for the residue, *W* and *b* is the weight matrix and bias learned by models and we choose ReLU (Glorot *et al.*, 2011) as the activation function *σ*.

### 2.5 Performance evaluation and measures

To evaluate the performance of models in this paper, we employed the following performance measures:

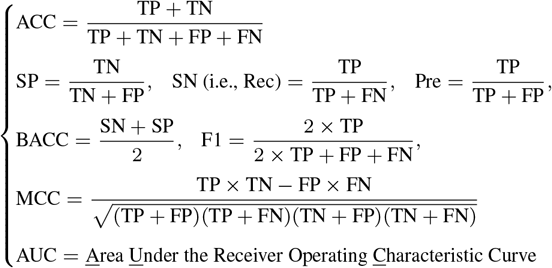

where TP, FP, FN, TN represent the true positive, false positive, false negative, and true negative respectively.

Note that AUC measure is quite different from others. In general, the predictors do not directly predict but give probabilities *p* for each residue. A threshold *δ* is chosen and if *p* ≤ *δ* then the prediction is 0 and otherwise 1. AUC does not rely on the choice of threshold *δ* while others do.

## 3 Results and discussion

The IDP-PLM is based on several related predictors for different tasks, including secondary structure prediction, protein-binding prediction, DNA-binding prediction, RNA-binding prediction, linker prediction and lipid-binding prediction. We describe and discuss the results for each of them and finally the IDP-PLM. The ablation analysis was also performed.

All predictors was implemented with Python 3.8.8 under PyTorch 1.11.0 framework (Paszke *et al.*, 2019) and CUDA 11.3 (NVIDIA *et al.*, 2020) on a PC with NVIDIA GeForce RTX 3090Ti GPU. Since the pre-trained ESM-1b model in our experiments was frozen during the training, we cached its outputs of all training sequences on the disk and reloaded them when training started, to avoid the repetitive redundant inferences of PLM across epochs. Consequently, both the GPU memory consumption and training time were significantly reduced. In the case of release, this trick would be removed and PLM inferences would be actually performed during the prediction.

### 3.1 Secondary structure prediction

Secondary structure (S.S.) prediction based on protein language models has been implemented and evaluated in quite a few works (Rao *et al.*, 2019; Rives *et al.*, 2021). Since there are no publicly available models, we implemented an S.S. predictor and its architecture is shown in Figure 1(b). Compared to the work (Rives *et al.*, 2021) which employed a convolutional layer and LSTM layers, our model adopted only a three-layer LSTM network after the PLM. Our model is trained by AdamW optimizer (Loshchilov and Hutter, 2017) with a learning rate of 0.0001, weight decay 0.01, 0.1 dropout rate and batch size of 64 under cross-entropy loss on the NetSurfP-2.0 dataset (Klausen *et al.*, 2019). We employed 5-fold cross-validation and the best weights were chosen regarding the maximum validated accuracy by employing the early stopping strategy to prevent the model from overfitting after stopping increasing the accuracy for more than 5 epochs. Meanwhile, the gradient clipping of value 1.0 was also adopted to avoid potential gradient explosions. The final result is the average accuracy on each fold.

We tested these cross-validated models in predicting 8 different (Q8) secondary structures on the CB513 independent dataset (Cuff and Barton, 1999) and compared the results with several state-of-the-art methods as shown in Table 1. Our predictor achieved the best performance in terms of accuracy compared to other PLM-based works. ProtT5-XL-U50 (Elnaggar *et al.*, 2020) is a PLM-based method that achieved the second-best performance by leveraging different pre-trained models from ESM-1b. Here ESM-1b-SS (Rives *et al.*, 2021) is the work associated with ESM-1b and its result is reported according to the original paper. In addition, DeepCNF-SS (Wang *et al.*, 2016b) and DeepSeqVec (Heinzinger *et al.*, 2019) are also two PLM-based methods that adopt relatively small-scale PLMs, and this also limited their performance.

**Table 1.** Comparison of Q8 secondary structure prediction on the CB513. Predictor Accuracy

| Predictor | Accuracy |
| --- | --- |
| **(This work)** | **75.1** |
| ProtT5-XL-U50 (Elnaggar *et al.*, 2020) | 74.5 |
| ESM-1b-SS (Rives *et al.*, 2021) | 70.2 |
| DeepCNF-SS (Wang *et al.*, 2016b) | 68.3 |
| DeepSeqVec (Heinzinger *et al.*, 2019) | 62.7 |

### 3.2 Linker prediction, protein-, DNA-, RNA-binding prediction, and lipid-binding prediction

We employed the same architecture as shown in Figure 1(b) to predict protein-binding, linker, DNA-binding, RNA-binding, and lipid-binding. These functional regions are relatively less occurred compared to the non-IDRs and this leads to the extremely imbalanced distributions between two different types of residues in the datasets. As a result, deep learning models are very sensitive and hard to converge on them.

We employed several training strategies to overcome the difficulties. A downsampling strategy is utilized to balance the large gap between the number of ordered and disordered residues in all datasets because the datasets for either annotated protein-, DNA-, RNA-binding sites (Zhang *et al.*, 2022), linkers (Meng and Kurgan, 2016) or lipid-binding sites (Katuwawala *et al.*, 2021) are extremely rare. For more details on our downsampling strategies, refer to the Supplementary Information. The focal loss (Lin *et al.*, 2017) was also employed to reduce the influence of imbalance in datasets. During the training, we adopted an AdamW optimizer and cross-validated the datasets. The early stopping strategy was also employed to prevent models from overfitting regarding the measure 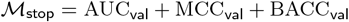 for specific epochs, since the single measure cannot indicate the performance comprehensively. For a more detailed explanation, refer to the Supplementary Information.

#### Protein-binding prediction

We trained and tested our protein-binding predictor on benchmark datasets in the reported work (Zhang *et al.*, 2022) and compared it with several state-of-the-art methods including DeepDISOBind (Zhang *et al.*, 2022) and DisoRDPbind (Oldfield *et al.*, 2020). The results of DisoRDPbind performance on the test dataset are evaluated by their official web service. As shown in Figure 2(a), our predictor achieved the best results on all measures including AUC, SN and F1, which demonstrates its high applicability. We conclude the advantages may come from PLMs because the other two used only traditional human-made features.

**Fig. 2.**
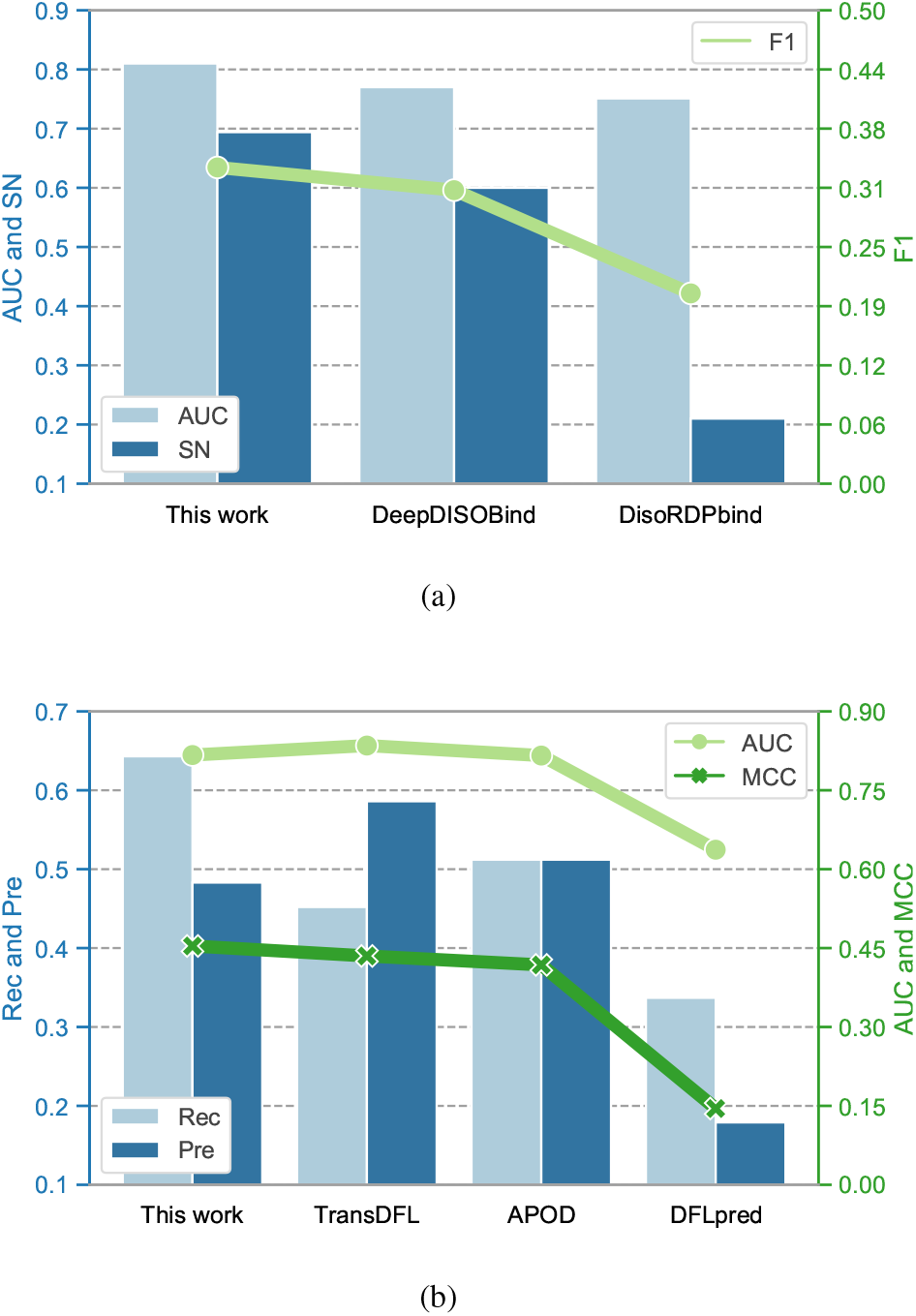
Comparison of different (a) protein-binding predictors and (b) linker predictors.

#### Linker prediction

We trained and tested our linker predictor on the benchmark datasets from the reported work (Meng and Kurgan, 2016) and compared it with several state-of-the-art methods including TransDFL (Pang and Liu, 2022), APOD (Peng *et al.*, 2020) and DFLpred (Meng and Kurgan, 2016). The results are shown in Figure 2(b). The final results are the average measures on each fold and our predictor achieved the best results on MCC, and Rec measures while having comparable results on AUC and Pre measure. TransDFL achieved the best performance on AUC and Pre measures and this is reasonable because it employed transfer learning by exploiting the knowledge from IDP prediction, while our linker predictor is intentionally trained for IDP prediction.

#### DNA- and RNA-binding prediction

We trained and tested our DNA- and RNA-binding predictors on benchmark from the reported work (Zhang *et al.*, 2022) and compared them with several state-of-the-art methods including DeepDISOBind (Zhang *et al.*, 2022) and DisoRDPbind (Oldfield *et al.*, 2020). Both of our two predictors achieved very high improvements in AUC and SN measures. However, the F1 measures are relatively lower than previous work. Since the F1 measure is the harmonic mean of Rec and Pre (see Section 2.5) and considering the high AUC measure, we conclude that the default threshold is not suitable for prediction. Despite this, it will not affect the following IDP prediction.

#### Lipid-binding prediction

We trained and tested our lipid-binding predictor on the benchmark from the reported work (Katuwawala *et al.*, 2021) and compared it with several state-of-the-art methods including DisoLipPred (Katuwawala *et al.*, 2021), ESpritz-DisProt (Walsh *et al.*, 2012) and SPOT-Disorder (Hanson *et al.*, 2017). The results are shown in Table 3. Our predictor achieved the best AUC measure and comparable results on SN and F1 measures. Despite DisoLipPred achieving better performance on SN and F1 measures, our predictor with the highest AUC measure suggests a better threshold is possible to adopt to improve SN and F1 measures. However, it will not influence the following IDP prediction.

**Table 2.** Comparison of DNA- and RNA-binding prediction on the test dataset.

| Type | Predictor | AUC | SN | F1 |
| --- | --- | --- | --- | --- |
| DNA | (This work) | **0.819** | **0.662** | 0.159 |
|  | DeepDISOBind (Zhang *et al.*, 2022) | 0.736 | 0.472 | **0.255** |
|  | DisoRDPbind (Oldfield *et al.*, 2020) | 0.671 | 0.452 | 0.246 |
| RNA | (This work) | **0.886** | **0.843** | 0.071 |
|  | DeepDISOBind (Zhang *et al.*, 2022) | 0.746 | 0.611 | **0.320** |
|  | DisoRDPbind (Oldfield *et al.*, 2020) | 0.594 | 0.364 | 0.202 |

**Table 3.** Comparison of lipid-binding prediction on the test dataset.

| Predictor | AUC | SN | F1 |
| --- | --- | --- | --- |
| (This work) | **0.817** | 0.377 | 0.093 |
| DisoLipPred (Katuwawala *et al.*, 2021) | 0.781 | **0.382** | **0.145** |
| ESpritz-DisProt (Walsh *et al.*, 2012) | 0.768 | 0.355 | 0.135 |
| SPOT-Disorder (Hanson *et al.*, 2017) | 0.692 | 0.155 | 0.062 |

### 3.3 Intrinsically disordered protein prediction

#### 3.3.1 Test on independent datasets

We tested our predictor on independent datasets DISORDER723, MXD494 and CAID. For fair comparisons, we removed the sequences in training datasets with identities greater than 25% from the test dataset and retrained the model from scratch. The distributions of the length of disordered regions are also shown in Supplementary Fig. S1 and the statistics of these datasets are shown in Supplementary Table S2. In addition, the DISORDER723 dataset has more short disordered regions while the CAID dataset has more long disordered regions and the MXD494 dataset has a balance between the other two. The results on above test datasets are shown in Tables 4, 5 and 6 where the previously reported results on DISORDER723 and MXD494 are taken from (Tang *et al.*, 2020) and CAID are taken from (Papastratis, 2022).

**Table 4.** Comparison of various IDP predictors on independent dataset DISORDER723.

|  | MCC | AUC | BACC | SN | SP |
| --- | --- | --- | --- | --- | --- |
| **IDP-PLM (This work)** | **0.623** | **0.939** | 0.795 | 0.609 | 0.981 |
| AUCpreD (Wang *et al.*, 2016a) | 0.564 | 0.914 | 0.777 | 0.580 | 0.974 |
| ESM1-t6-85M (Papastratis, 2022) | 0.548 | 0.769 | 0.770 | 0.565 | 0.973 |
| DISOPRED3 (Jones and Cozzetto, 2015) | 0.536 | 0.899 | 0.719 | 0.452 | 0.986 |
| IDP-Seq2Seq (Tang *et al.*, 2020) | 0.511 | 0.906 | 0.787 | 0.618 | 0.955 |
| RFPR-IDP (Liu *et al.*, 2021) | 0.442 | 0.898 | 0.748 | 0.522 | 0.974 |
| SPINE-D (Zhang *et al.*, 2012) | 0.376 | 0.891 | **0.810** | 0.779 | 0.840 |

**Table 5.** Comparison of various IDP predictors on independent dataset MXD494.

|  | MCC | AUC | BACC | SN | SP |
| --- | --- | --- | --- | --- | --- |
| **IDP-PLM (This work)** | **0.485** | **0.851** | 0.720 | 0.515 | 0.925 |
| IDP-Seq2Seq (Tang *et al.*, 2020) | 0.475 | 0.825 | **0.767** | 0.782 | 0.756 |
| ESM1-t6-85M (Papastratis, 2022) | 0.467 | 0.749 | 0.743 | 0.633 | 0.854 |
| SPOT-Disorder (Hanson *et al.*, 2017) | 0.457 | 0.813 | 0.739 | 0.626 | 0.851 |
| RFPR-IDP (Liu *et al.*, 2021) | 0.442 | 0.821 | 0.754 | 0.749 | 0.758 |
| SPINE-D (Zhang *et al.*, 2012) | 0.411 | 0.803 | 0.742 | 0.787 | 0.698 |
| AUCpreD (Wang *et al.*, 2016a) | 0.411 | 0.800 | 0.701 | 0.512 | 0.881 |

**Table 6.** Comparison of various IDP predictors on independent dataset CAID.

|  | MCC | AUC | F1 | BACC | SN | SP |
| --- | --- | --- | --- | --- | --- | --- |
| **IDP-PLM (This work)** | 0.369 | **0.814** | **0.483** | 0.717 | 0.617 | 0.817 |
| fIDPnn (Hu *et al.*, 2021) | **0.370** | **0.814** | **0.483** | **0.720** | 0.629 | 0.811 |
| ESM1-t6-85M (Papastratis, 2022) | 0.365 | - | 0.460 | 0.715 | 0.691 | 0.772 |
| fIDPlr (Hu *et al.*, 2021) | 0.330 | 0.793 | 0.452 | 0.693 | 0.570 | 0.816 |
| SPOT-Disorder (Hanson *et al.*, 2017) | 0.330 | - | 0.434 | 0.723 | 0.832 | 0.614 |
| AUCpreD (Wang *et al.*, 2016a) | 0.318 | - | 0.433 | 0.712 | 0.801 | 0.624 |

We adopted the 5-fold cross-validation and choose the best model regarding the aforementioned stop metric 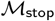. Moreover, the early stopping strategy was also employed to prevent overfitting when it stop increasing for 3 epochs. We adopted an AdamW optimizer with a learning rate of 0.0005, weight decay of 0.01 and batch size of 32.

##### DISORDER723

IDP-PLM significantly improved the performance in terms of AUC and MCC measures on DISORDER723, as shown in (Table 4). It has the second-best BACC measure compared with the SPINE-D. However, SPINE-D has lower AUC and MCC than ours. We note that the recent work to ESM1-t6-85M (Papastratis, 2022), which is also based on PLMs and ESM, achieved relatively lower measures than ours.

##### MXD494

IDP-PLM significantly outperforms in terms of MCC and AUC measures on the dataset MXD494 while slightly lower than IDP-Seq2Seq (Tang *et al.*, 2020) on BACC, as shown in (Table 5). However, IDP-Seq2Seq has a lower AUC and MCC than ours. Since the AUC measure is more comprehensive and the BACC measure is sensitive to the thresholds chosen by users, our predictor is more robust in practice.

##### CAID

IDP-PLM achieved state-of-the-art performance in terms of AUC and F1 measures and a very close MCC and BACC measures compared to the latest work (Hu *et al.*, 2021) on CAID (since the AUC measures are not available for all predictors, we also listed the F1 measure), as shown in (Table 6). This is because CAID is very difficult to predict compared to other datasets such as DISORDER723 and MXD494. We note that flDPnn achieved the best performance because it used various high-quality input profiles including MSA-based features. This also suggests that the MSA-based features are more important than representations extracted from PLMs.

We conclude that the improvements of our method may come from 1) the good representations extracted by PLM, which may contain essential information about proteins, and 2) our well-designed stacked architecture and BiLSTM-based predictors greatly contributed. In contrast, ESM1-t6-85M used a simple soft-max layer appending the last layer of the PLM, which could not exploit the sequential information from the PLM effectively. 3) we employed a larger training dataset. All of these make IDP-PLM a more powerful predictor.

#### 3.3.2 Ablation study

To determine the contributions of each module in our model, we performed the ablation study by removing the specific module, retraining the model from scratch and testing it on the CAID dataset because it is the most challenging dataset. As described in Section 2, the full version of IDP-PLM consists of seven major modules including a secondary structure predictor, a protein-binding predictor, a linker predictor, a DNA-binding predictor, an RNA-binding predictor, a lipid-binding predictor and an ADD-MEAN module. To observe the effect of each module, we tested the performance on the CAID dataset by separately removing them, i.e., the protein-binding predictor (-Prot.), the DNA-binding predictor (-DNA), the RNA-binding predictor (-RNA), linker predictor (-Linker), the lipid-binding predictor (-Lipid), the secondary structure predictor (-S.S.) and ADD-MEAN module (-A.M.). The results are shown in Figure 3 and Supplementary Information Table S1. All modules in the IDR predictor contribute positively to the performance of the IDP-PLM. Compared to other modules, removing the lipid-binding, secondary structure predictor and ADD-MEAN module, however, brings a slightly larger decrease on all measures. This suggests that they are more informative and challenging to predict from the sequence. On the other hand, the linker predictor contributes the least to the overall performance. This is because of the relatively small scale of linker datasets, even though it performed well on the test datasets.

**Fig. 3.**
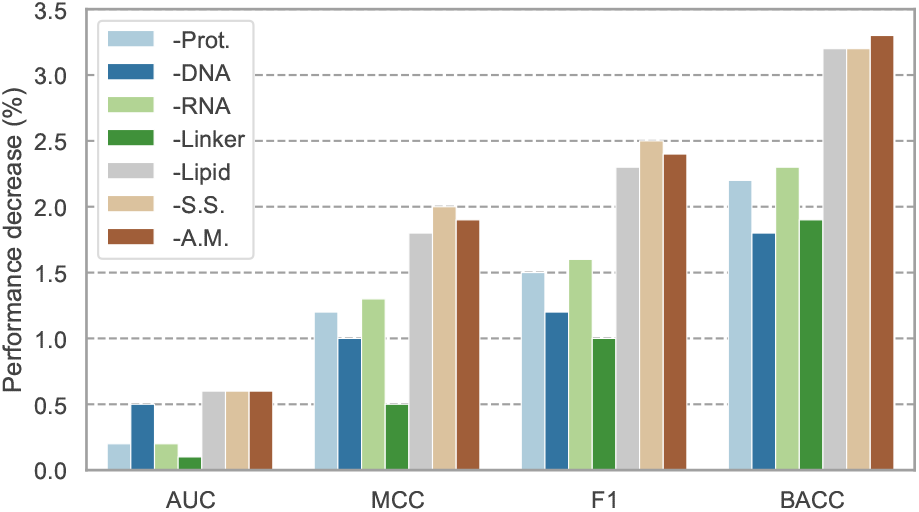
Ablation study on the independent dataset CAID.

## 4 Conclusion

We proposed a novel, accurate yet efficient IDR prediction method based on the protein language model (PLM) and achieved state-of-the-art performance. Meanwhile, we developed several independent predictors which can be used separately for linker prediction, protein-, DNA-, RNA-binding prediction, and lipid-binding prediction. These predictors achieved state-of-the-art results in terms of accuracy and efficiency. Our work demonstrates that architecture based on PLM and utilizing traditional input features can improve the performance of IDR prediction. In the future, we hope to extend the IDR prediction with more input features such as the predicted residue-residue contact information extracted from PLM attention maps.

