## Supporting Information for "Fast and Accurate Prediction of Intrinsically Disordered Protein by Protein Language Model"

### Supplementary Information for Fast and Accurate Prediction of Intrinsically Disordered Protein by Protein Language Model

#### 1. FIGURES AND TABLES

**Table S1.** Statistics of test datasets for protein-, DNA-, RNA-binding, linker and lipid-binding sites. The number of binding sites or linker residues (#Pos. resi.), the number of other residues (#Neg. resi.) and the number of proteins (#Prot.) for each dataset are listed.

| | Task | #Pos. resi. | #Neg. resi. | #Prot. |
|---|---|---|---|---|
| Train | Linker | 2003 (8.7%) | 21151 (91.3%) | 148 |
| | Protein-binding | 33736 (10.4%) | 290137 (89.6%) | 767 |
| | DNA-binding | 5611 (1.7%) | 318262 (98.3%) | 767 |
| | RNA-binding | 3284 (1.0%) | 320589 (99.0%) | 767 |
| | Lipid-binding | 1921 (2.0%) | 94094 (98.0%) | 222 |
| Test | Linker | 922 (9.0%) | 9294 (91.0%) | 61 |
| | Protein-binding | 4608 (13.9%) | 28639 (86.1%) | 86 |
| | DNA-binding | 963 (2.9%) | 32284 (97.1%) | 86 |
| | RNA-binding | 279 (0.8%) | 32968 (99.2%) | 86 |
| | Lipid-binding | 1471 (1.4%) | 104877 (98.6%) | 252 |

**Table S2.** Statistics of independent datasets. The number of disordered residues (#Dis. resi.), the number of ordered residues (#Ord. resi.) and the number of proteins (#Prot.) for each dataset are listed.

| | Datasets | #Dis. resi. | #Ord. resi. | #Prot. |
|---|---|---|---|---|
| Train | DISORDER723* | 435426 (12.2%) | 3124798 (87.8%) | 11541 |
| | MXD494* | 392537 (11.2%) | 3126472 (88.8%) | 11556 |
| | CAID* | 326009 (10.6%) | 2757572 (89.4%) | 10769 |
| Test | DISORDER723 | 13490 (6.3%) | 201228 (93.7%) | 726 |
| | MXD494 | 43475 (22.2%) | 152414 (77.8%) | 517 |
| | CAID | 54825 (16.3%) | 282081 (83.7%) | 731 |

mark "*" means the full training dataset removing redundant homologous sequences from the test datasets.

**Table S3.** Ablation study on the independent CAID dataset.

| Model | AUC | MCC | F1 | BACC | SN (i.e., Rec) | SP | Pre |
|---|---|---|---|---|---|---|---|
| Full | 0.814±0.000 | 0.369±0.003 | 0.483±0.002 | 0.717±0.002 | 0.617±0.017 | 0.817±0.012 | 0.398±0.010 |
| - Linker | 0.812±0.002 | 0.357±0.005 | 0.468±0.004 | 0.695±0.007 | 0.537±0.035 | 0.853±0.021 | 0.424±0.019 |
| - Prot. | 0.809±0.002 | 0.359±0.005 | 0.471±0.004 | 0.699±0.007 | 0.551±0.033 | 0.847±0.020 | 0.418±0.017 |
| - A.M. | 0.812±0.002 | 0.356±0.006 | 0.467±0.007 | 0.694±0.009 | 0.532±0.034 | 0.856±0.017 | 0.423±0.012 |
| - S.S. | 0.813±0.001 | 0.364±0.002 | 0.473±0.001 | 0.698±0.004 | 0.629±0.019 | 0.810±0.011 | 0.393±0.007 |
| - LIPID | 0.808±0.002 | 0.351±0.005 | 0.460±0.002 | 0.685±0.006 | 0.501±0.037 | 0.868±0.025 | 0.436±0.021 |
| - DNA | 0.808±0.002 | 0.350±0.005 | 0.459±0.002 | 0.684±0.006 | 0.500±0.036 | 0.868±0.024 | 0.436±0.021 |
| - RNA | 0.808±0.002 | 0.349±0.004 | 0.458±0.002 | 0.685±0.007 | 0.506±0.039 | 0.863±0.026 | 0.430±0.021 |

**Table S4.** Hyper-parameters for linker predictor, protein-, DNA-, RNA-, lipid-binding predictor, secondary structure predictor and IDP-PLM.

| Predictor | number of fold | learning rate | weight decay | batch size | mask ratio | patience |
|---|---|---|---|---|---|---|
| Linker | 10 | $10^{-3}$ | $10^{-5}$ | 64 | 0.05 | 30 |
| Protein-binding | 10 | $10^{-3}$ | $10^{-5}$ | 64 | 0.05 | 30 |
| DNA-binding | 5 | $10^{-4}$ | $10^{-5}$ | 16 | 0.001 | 20 |
| RNA-binding | 5 | $10^{-4}$ | $10^{-5}$ | 16 | 0.001 | 20 |
| Lipid-binding | 4 | $5 \cdot 10^{-5}$ | $10^{-5}$ | 16 | 0.0005 | 20 |
| Secondary structure | 10 | $10^{-4}$ | $10^{-2}$ | 64 | - | 5 |
| IDP-PLM | 5 | $10^{-4}$ | $10^{-2}$ | 32 | - | 3 |

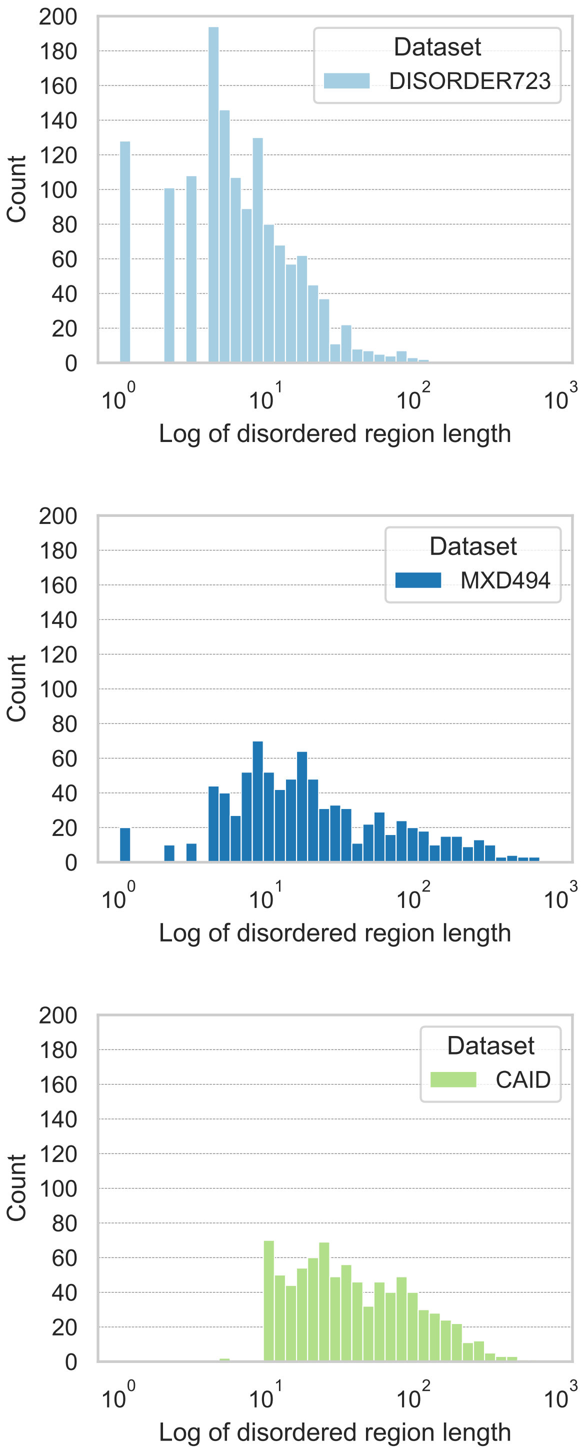


**Fig. S1.** Distributions of the averaged length of disordered regions contained in DISORDER723, MXD494 and CAID. Log-axes are enabled for x-axis.

#### DOWNSAMPLING STRATEGIES

We employed an efficient downsampling method by selecting unmasked residues according to one of the following conditions:

- each residue is selected with a specific probability *delta* (which named "mask ratio" in aforementioned context),
- the residue is annotated as positive (i.e., it is a protein-binding residue or a linker).

To be more specific, the pseudo-code is shown as follows:

```
def train(batch):
    inputs, targets = batch
    # set up a mask
    mask = ((torch.rand(targets.shape) < delta ) | (targets != 0))
    # forward propagation
    y_pred = model(inputs)
    loss = loss_func(y_pred[mask], targets[mask])
    # ....
```
